# Massive accumulation of strontium and barium in diplonemid protists

**DOI:** 10.1101/2022.06.27.497835

**Authors:** Jana Pilátová, Daria Tashyreva, Jiří Týč, Marie Vancová, Syed Nadeem Hussain Bokhari, Radim Skoupý, Mariana Klementová, Hendrik Küpper, Peter Mojzeš, Julius Lukeš

## Abstract

Barium (Ba) and strontium (Sr) are often used as proxies for the reconstruction of past marine productivity and global climate. The ability to accumulate Ba^2+^ and Sr^2+^ in the form of crystals is rare among eukaryotes. Here we report that unicellular heterotrophs called diplonemids (Euglenozoa), one of the most abundant groups of marine planktonic protists, accumulate conspicuous amounts of these trace elements in the form of intracellular barite (BaSO_4_) and celestite (SrSO_4_) crystals, in concentrations greater than in other known Ba/Sr-accumulating organisms. Moreover, these flagellates can uptake Sr^2+^ exclusively or together with Ba^2+^ and form (Ba,Sr)SO_4_. One species, *Namystinia karyoxenos*, is naturally capable of intracellular accumulation of Ba^2+^ and Sr^2+^ 42,000 and 10,000 times relative to the surrounding medium. Altering the amounts of both elements in the medium resulted in corresponding changes in the quantity and composition of crystals. Planktonic copepods fed with diplonemids produce celestite-rich fecal pellets, which facilitate deposition of these minerals on the seafloor. We propose that diplonemids, which emerged during the Neoproterozoic era, qualify as impactful players of Ba^2+^/Sr^2+^ cycling in the ocean that possibly contributed to sedimentary rock formation over long geological periods.

## INTRODUCTION

Although in most environments strontium (Sr) and barium (Ba) are present in trace amounts, they can be accumulated in substantial quantities by some organisms [1, 2]. Depending on their environmental availability, these elements are mostly taken up non-selectively together with Ca^2+^ [3, 4]. The soluble form of Ba^2+^ is typically toxic for animals (*e.g*., used as rodenticides) due to its capacity to block K^+^ channels, while insoluble BaSO_4_ acts as a common contrast agent in medical radio-imaging [5]. In contrast, soluble Sr^2+^ is not harmful, with the exception of the radioactive isotope ^90^Sr^2+^ occurring as nuclear contaminants that accumulate in marine biota and sediments [6]. Indeed, in some algae, Sr^2+^ can almost fully replace Ca^2+^ without any discernable deleterious effects [7]. In humans, Sr^2+^ treatment of osteoporosis is used to prevent fractures [8]. Moreover, predictions concerning climate change stress the increased relevance of higher environmental mobilization of Sr^2+^ and Ba^2+^ due to enhanced solubility upon marine acidification [9]. Apart from chemical precipitation treatments of radioactive ^90^Sr^2+^ and toxic Ba^2+^ there are new attempts for bioremediation using cyanobacteria, algae and fungi [1, 2, 6, 10].

In marine environments, microorganisms accumulate more Sr^2+^ than Ba^2+^ possibly due to higher solubility and availability of the former element [9]. In protists, Sr^2+^ is mostly present in the form of celestite (also referred to as celestine; SrSO_4_) and strontianite (SrCO_3_), while Ba^2+^ forms barite (BaSO_4_) or witherite (BaCO_3_). Moreover, Ba^2+^ and Sr^2+^ commonly substitute each other in various ratios to form strontiobarite and baritocelestite (Ba,Sr)SO_4_ [11]. Celestite with traces of Ba^2+^ is well known for forming the complex skeletons of acanthareans [12]. Intracellular barite crystals form statoliths of some freshwater charophyte algae and statocysts of marine ciliates, in which they likely play a role in graviperception [13–15]. Haptophytes and foraminiferans form intracellular barite crystals with trace amounts of Sr^2+^ [16, 17], while strontianite and witherite occur in microalga *Tetraselmis* [18] and coccolithophorids [19, 20]. The exact role of these crystalline inclusions remains unknown.

Marine Ba^2+^ and Sr^2+^ are frequently correlated with particulate organic carbon in the water column and sediments on the sea floor, indicating that microorganisms are capable of accumulating these elements [21–23], yet the celestite-rich skeletons of acanthareans dissolve during sedimentation [24]. Ba^2+^ and Sr^2+^ carbonates and phosphates known from coccolithophorids and bacteria, respectively, contribute to the cycling of these elements with possible conversion to sulfates in the process of diagenesis [10, 19, 20, 25]. In addition, barite and strontiobarite crystals were frequently found associated with fecal pellets, which contribute to sedimentation of particulate Ba^2+^ and Sr^2+^ to the sea floor [23]. However, until now, abundant planktonic organisms capable of selective intracellular accumulation of both Ba^2+^ and Sr^2+^ sulfates have not been identified [15, 26]. Despite the well-documented evidence-based predictions of the biogenic origin of barite and celestite minerals in the oceans [23, 27], the lack of organisms responsible for their production led to the gradual focus on microenvironment-mediated precipitation – stepping away from consideration of their biological origin [28].

Here, we show that diplonemids (Diplonemea, Euglenozoa), a group of biflagellated heterotrophic protists [29–31], are capable of massive intracellular accumulation of Sr^2+^ and Ba^2+^. Specifically, three cultivable diplonemids accumulate celestite and barite crystals in intracellular concentration of Sr^2+^ much greater than in other organisms [10, 16]. Diplonemids have only recently been recognized as one of the most diverse and abundant groups of microeukaryotes in the world ocean, especially in the mesopelagic zone [32–34]. Although relatively rare, they are present in freshwater bodies as well [35]. We analyze their crystalline inclusions by a range of complementary approaches and discuss their possible biological functions and role in biogeochemical cycles.

## MATERIALS AND METHODS

### Cell cultures, cultivation and light microscopy

For all experiments, axenic cultures were grown in seawater-based Hemi medium (SI Appendix Table S1) supplemented with 1% horse serum and 0.025 g/L LB broth powder [36]. An artificial seawater medium lacking Sr^2+^, Ba^2+^ and sulfates was prepared from 288 mM NaCl, 8 mM KCl, 718 mM KBr, 100 mM MgCl_2_; 12 mM CaCl_2_; 40 mM HBO_3_, 60 mM NaF, supplemented with 1% (v/v) heat-inactivated horse serum (Sigma-Aldrich) and 25 mg LB broth powder (Amresco). The medium was used as rinsing solution for preparation of ICP-MS samples. For Ba^2+^ loading experiments, this medium was supplemented with BaCl_2_ equimolarly to the naturally occurring Sr^2+^ of 8 mg·l^−1^.

Axenic clonal cultures of 21 strains of diplonemids were grown either at 27 °C (*Paradiplonema papillatum* ATCC 50162), 22 °C (*Namystinia karyoxenos* YPF1621) or 13 °C (of *D. aggregatum* YPF1605, *D. japonicum* YPF1604, *Flectonema* sp. DT1601, *Hemistasia phaeocysticola, Lacrimia lanifica* JW1601 and *Lacrimia* sp. YPF1808, *Rhynchopus* sp. YZ270 cl. 10.3, *Rhynchopus* sp. YZ270, cl. 9, *Rhynchopus* sp. DT0301, *Rhynchopus humris* (strains YPF1505, FC902, FC904, and KQ12), *R. euleeides* (strains YPF1915 and ATCC 50226), *R. serpens* YPF1515, *Sulcionema specki* YPF1618). The identity of not yet formally described species was established based on the 18S rRNA sequences as described previously [37]. Dense cultures of trophic cells [38] were harvested by centrifugation at 3,000× g for all subsequent analyses.

Light microscopy images and videos were taken with an Olympus BX53 microscopy equipped with DP72 microscope digital camera using CellSens software v. 1.11 (Olympus), and processed with GIMP v. 2.10.14, Irfan View v. 4.54 and Image J v. 1.51 software. Polarized microscopy was performed using crossed polarizers installed to Raman microscope (as specified below).

### Environmental sampling

Zooplankton was collected in the Bay of Villefranche sur Mer (France: 43°40’N, 7°19’E) with a 10 min haul from 10 m to the surface, using a 20 μm mesh size plankton-net. Captured copepods were transferred into 0.5 l of freshly filtered natural seawater and starved for 12 hrs. *Centropages typicus*, *Temora longicornis* and *Acartia* sp. were then picked under a dissection microscope. All experiments were carried out at cultivation temperature of the prey species of diplonemids (13°C for *Lacrimia* sp. YPF1808, room temperature for *N. karyoxenos*). Ten copepods were kept in 20 ml of diplonemid culture (10^5^ cells ml^−1^) for 5 days, after which their fecal pellets were collected under a dissection microscope and immediately analyzed by Raman microscopy (as specified below).

### Raman microscopy

For the *in situ* determination of the chemical composition of intracellular structures, a confocal Raman microscope WITec alpha300 RSA (WITec, Germany) was used as previously described in [39–42]. To immobilize the fast-moving flagellates on the quartz slide, 5 μl of the cell pellet was mixed with 5 μl of 1% w/v solution of low-melting agarose (Art. Nr. 6351.5, Carl Roth, Germany), immediately spread as a single-cell layer between a quartz slide and coverslip and sealed with a CoverGrip sealant (Biotium, USA). Two-dimensional Raman maps were obtained with a laser excitation at 532 nm (20 mW power at the focal plane) and oil-immersion objective UPlanFLN 100×, NA 1.30 or water-immersion objective UPlanSApo 60×, NA 1.20 (Olympus, Japan). A scanning step size of 200 nm in both directions, and an integration time of 100 ms per voxel were used. A minimum of 30 cells were measured for each strain. Raman chemical maps were constructed by multivariate decomposition of the baseline-corrected spectra into the spectra of pure chemical components by using WITec Project Plus 5.1 software (WITec, Germany).

### Transmission electron microscopy (TEM), electron diffraction (TEM-ED) and energy dispersive X-ray spectroscopy (TEM-EDX)

The protocol for the basic sample preparation of all kinds of electron microscopy approaches listed here is described in detail in [43]. We used it with minor modifications, as stated below. Cell pellets were transferred to specimen carriers and immediately frozen in the presence of 20% w/v bovine serum albumin (BSA) solution using a high-pressure freezer Leica EM ICE (Leica Microsystems, Austria). Freeze substitution was performed in the presence of 2% osmium tetroxide diluted in 100% acetone at −90 °C. After 96 h, specimens were warmed to – 20 °C at a step of 5 °C/h. After another 24 h, the temperature was increased to 3 °C (3 °C/h). At room temperature, samples were washed in acetone and infiltrated with 25%, 50%, 75% acetone/resin mixture for 1 h at each step. Finally, samples were infiltrated in 100% resin and polymerized at 60 °C for 48 h. Semi-thin (250 nm) and ultrathin (70 nm) sections were cut using a diamond knife, placed on copper grids, and stained with uranyl acetate and lead citrate. TEM micrographs were taken with Mega View III camera (SIS) using a JEOL 1010 TEM operating at an accelerating voltage of 80 kV.

For TEM-EDX, 10 μl of pelleted *L. lanifica* cells were spread over a holey carbon-coated copper grid, washed twice with 10 μl of distilled water in order to reduce the sea salts from the culture medium, and allowed to dry by evaporation at ambient temperature. Semi-thin sections of resin-infiltrated blocks of *N. karyoxenos* were prepared as stated above. For the identification of the crystalline phase, sections were studied by TEM on FEI Tecnai 20 (LaB6, 120 kV) equipped with an Olympus SIS CCD camera Veleta (2048 × 2048 px), and an EDAX windowless EDX detector Apollo XLTW for elemental analysis. The diffraction data were collected by the means of three-dimensional electron diffraction (3D ED) [44]. The data processing was carried out using the PETS software [45]. Structure solution and refinement were performed in the computing system Jana2006 [46].

### Cryo-scanning electron microscopy with energy dispersive X-ray spectroscopy (cryo-SEM-EDX)

Cells pellets were high-pressure frozen as described above and transferred into a Leica ACE 600 preparation chamber (Leica Microsystems, Austria) precooled at −135 °C, fractured with a scalpel, freeze-etched at −100 °C for 1 min, and sputter-coated with 2.5 nm of gold-palladium at −125 °C. Specimens were transferred under vacuum using a transfer system VCT100 (Leica Microsystems, Austria) and observed by the scanning electron microscope (SEM) Magellan 400L (FEI, Czech Republic/USA) pre-cooled at −125 °C (cryo-SEM). Topographical images and EDX measurements were taken using EDT detector and EDAX detector (Octane Elect Super; EDAX, USA), respectively, either at 5 keV, 0.1 nA, or 10 keV, 0.8 nA. The taken spectra were analyzed in EDAX TEAM software and quantified by eZAF method.

### Serial block-face scanning electron microscopy (SBF-SEM)

The sample preparation of *Lacrimia* sp. YPF1808 by high-pressure freezing technique follows the protocol for TEM sample preparation. After freeze-substitution, the samples were subsequently stained with 1% thiocarbohydrazide in 100% acetone for 1.5 hours, 2% OsO_4_ in 100% acetone for 2 hours at room temperature, and 1% uranyl acetate in 100% acetone overnight at 4°C. After every staining step, the samples were washed 3 times with 100% acetone for 15 min. Samples were then infiltrated with 25%, 50%, 75% acetone/resin mixture for 2 hours at each step, and finally infiltrated in 100% Hard Resin Plus 812 (EMS) overnight and polymerized at 62°C for 48 hours. Resin-embedded blocks were trimmed and imaged using Apreo SEM equipped with the VolumeScope (Thermo Fisher Scientific, Germany). Serial images were acquired at 3.5 keV, 50 pA, 40 Pa with a resolution of 6 nm and 100 nm slice thickness and dwell time per pixel of 4 μs. Image data was processed in Microscopy Image Browser v2.702 [47] and Amira v2020.2. The resin-embedded blocks were also collected in the form of 1 μm thick sections on the silicon wafer and analyzed by SEM-EDX (Magellan 400L as described above).

Based on volumetric data, we calculated the percentage of increase in cell density based on measured volumes of crystals compared to the theoretical crystal-free cells of the same volume and reported average theoretical density of 1.07 g·cm^−3^ [48].

### Inductively coupled plasma mass spectrometry (ICP-MS)

For analysis of Ba and ^88^Sr concentration, cultures were grown in triplicates, counted, and washed three times with 1 M sorbitol solution (*D. papillatum*, *D. japonicum* and *Rhynchopus* YZ270 cl. 10) or Sr- and Ba-free artificial seawater rinsing solution (see above) (*L. lanifica* JW1601, *Lacrimia* sp. 1808 and *N. karyoxenos*) to remove Ba^2+^ and Sr^2+^ present in the cultivation medium. Cultivated cells were harvested by centrifugation, each rinsed twice with 50 ml and once with 2 ml of the rinsing solution, and the resulting pellets were freeze-dried. A half ml of digestion acid mix (425 μl of 70% HClO_4_ and 75 μl of 69% HNO_3_) prepared as in [49] was added directly to the dried biomass. The digestion was done using a Fuji PXG4 Thermoblock (AHF Analysentechnik AG, Germany). After evaporation of the acid mix, 0.5 ml of 5% HCl was added to each test tube to re-dissolve the salts. The glass tubes were heated to 90 °C for 1 h to obtain clear solutions. The final volume of 1.5 ml was adjusted with ddH2O. Appropriate dilutions were done with 0.2% HNO_3_. Indium was added as an internal standard at 1 ng/ml to each test solution. The ICP multi-element standard solution VI (Merck, Germany) was used to prepare standard curves. Analyses were done using the inductively coupled plasma sector-field mass spectrometer (ICP sfMS) Element XR-2 with jet interface (Thermo Fisher Scientific, Germany) following a described protocol [50]. Medium resolution of 4000 was used in Ba and ^88^Sr measurements in triplicate of each technical replicate, with the highest precision and lowest relative standard deviation. Additionally, the elemental composition of samples of standard growth medium and artificial seawater medium without sulfates, Ba^2+^ and Sr^2+^ was analyzed.

### Holographic microscopy – quantitative phase imaging (QPI)

Samples for holographic microscopy were immobilized prior to measurement as described in the case of Raman microscopy. Imaging was performed at Q-Phase microscope (Tescan Orsay Holding, Czech Republic). Holographic Q-Phase microscope (Tescan Orsay Holding, Czech Republic) is equipped by halogen lamp illumination through the interference filter (λ ¼ 650 nm, 10 nm FWHM) and microscope objective (Nikon Plan Fluor oil immersion 60× NA 1.4, providing lateral resolution of 0.57 μm). The numerical reconstruction of acquired data was performed using Q-Phase software (Tescan Orsay Holding, Czech Republic). The technique enables automated cell segmentation and quantitative analysis of cellular mass based on the specific proportions of thickness and refractive indices of measured cells in comparison to the reference [51]. Due to the high variability of cell contents and sizes, at least 150 cells were analyzed for each strain. Because crystalline inclusions caused artifacts during capturing due to the big difference in refractive indices, we analyzed crystal-free cells cultivated in the artificial seawater medium lacking Sr^2+^, Ba^2+^ and sulfates (specified above). We calculated the total dry mass of the cells as sum of crystal-free cells measured by holographic microscopy and SrSO_4_ and BaSO_4_ amounts measured via ICP-MS. The dry-weight ratios of trace elements measured via ICP-MS were calculated based on the total dry weight of corresponding strains.

### Statistical data analysis

Statistical analysis we conducted using SigmaPlot, v. 12.5 and SPSS v. 23.0. Logarithmically normalized data were subjected to statistical tests (one-way ANOVA and Tukey’s post-hoc) on the level alpha of 0.05. Calculations of standard error of the mean based on independent methods (*i.e*., ICP-MS quantification and QPI dry mass quantification) with different levels of variability were done according to mathematical conversion using Taylor expansion.

## RESULTS

### Light microscopy and analysis of crystals by Raman microscopy

To determine the chemical composition of biogenic crystals directly within intact cells, Raman microscopy, a vibrational spectroscopic method sensitive to molecular composition, was used. Out of 21 strains belonging to 15 diplonemid species, three members of the distantly related genera *Lacrimia* and *Namystinia*, represented by *Lacrimia* sp. YPF1808, *L. lanifica*, and *Namystinia karyoxenos* were shown to possess celestite crystals (Fig. 1A). Their Raman spectra were congruent with the spectra of mineral celestite and chemically prepared precipitates of SrSO_4_ (Fig. 1F), matching also Raman spectra of celestite reported elsewhere [52]. Due to the counter-cation sensitivity of the position of the most intense Raman band at around 1,000 cm^−1^ belonging to the symmetric ν1 vibrational mode of SO_4_^2-^ tetrahedron, biogenic celestite could be unambiguously identified as SrSO_4_, and was easily distinguishable from barite, baritocelestite, gypsum (CaSO_4_) or calcite (CaCO_3_) (SI Appendix Fig. S1). Small relative-intensity nuances of other Raman bands of biogenic celestite from various cells (SI Appendix Fig. S2) can be explained by differences in crystal structures (lattice defects), trace admixtures of Ba^2+^ and/or orientation of the crystals, as these are present also in the spectra of mineral reference and chemical precipitates.

**Figure 1:**
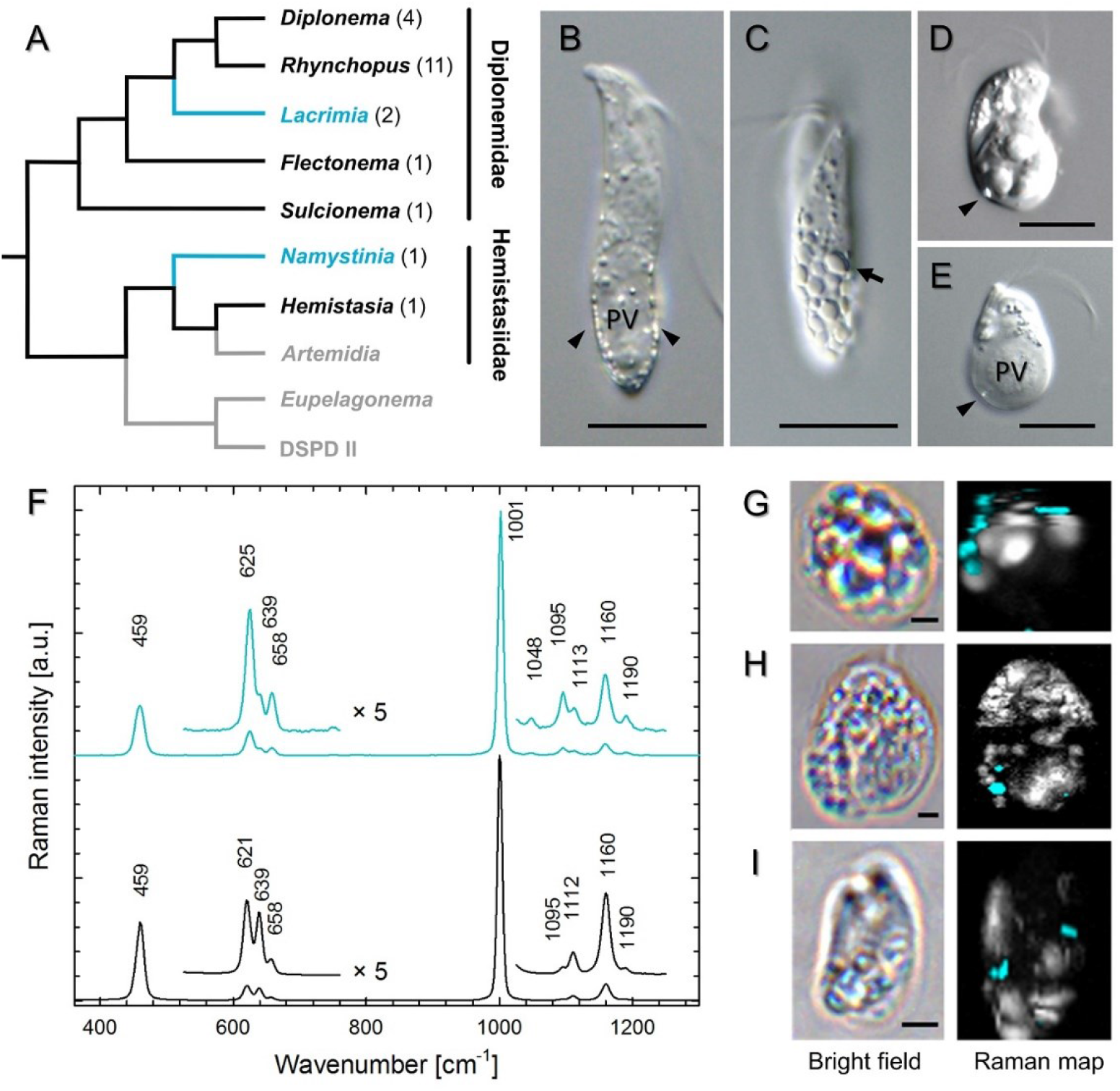
Distribution of celestite in diplonemids based on Raman microscopy analysis. (*A*) Phylogenetic tree of diplonemids based on [29] with genera containing celestite (blue), other screened genera (black) and not examined clades (grey). Differential interference contrast micrographs of *N. karyoxenos (B, C) Lacrimia* sp. YPF1808 (*D*) and *L. lanifica* (*E*) with celestite crystals marked by *arrowheads; arrow* points to large polygonal crystal. Scale bar = 10 μm. (*F*) Raman spectra of biogenic celestite crystals found in diplonemid cells (blue) and celestite mineral (black). Raman chemical maps of *N. karyoxenos* (*G*), *Lacrimia* sp. YPF1808 (*H*), and *L. lanifica* JW1601 (*I*) with celestite in blue and other cytoplasmic contents in white. Scale bar = 2 μm.

When *N. karyoxenos* was examined by light microscopy with differential interference contrast (DIC), the crystalline structures appeared as small birefringent particles moving fast by Brownian motion (Fig. 1B, C; SI Appendix Movie 1). They were most prominent within the enlarged lacunae, which are peripheral membrane-bounded compartments positioned directly beneath the subpellicular microtubular corset (SI Appendix Movie 1) following the addition of tiny amounts of formaldehyde. Within a single culture, the size and quantity of crystals inside the cells ranged from a few small particles (Fig. 1B) up to multiple large, tightly packed, polygonal crystals reflecting the shape of orthorhombic prisms (Fig. 1C). The crystalline particles of *Lacrimia* sp. YPF1808 and *L. lanifica* were far less prominent under the light microscope than those of *N. karyoxenos*. However, large crystals were visible around the posterior vacuole with DIC (Fig. 1D, E; SI Appendix Movie 2) and under polarized light (SI Appendix Movies 3 and 4).

### Morphology, localization and elemental analysis of intracellular crystals

Examination with light, Raman, transmission electron microscopy (TEM) and serial block face scanning electron microscopy (SBF-SEM), showed that the crystalline inclusions in two clades of diplonemids differed in their localization and shapes. In semi-thin resin-embedded sections of *N. karyoxenos*, numerous orthorhombic prismatic and bipyramidal crystals were localized mostly inside the lacunae (Fig. 2A, C, D), with a preference towards the cell posterior. Occasionally, crystals were found inside the large posterior vacuole (Fig. 2A) and in smaller vacuoles scattered throughout the cytoplasm (Fig. 2A, B). Only small crystals could be seen in semi-thin sections, while bigger crystals dropped out leaving empty crystal-shaped holes. Due to frequent rupturing, it was not possible to visualize celestite crystals in semi-thin epoxy resin sections of *Lacrimia* species. Thus, we used the SBF-SEM approach, which showed that the celestite crystals of *Lacrimia* sp. YPF1808 appeared mostly in small membrane-bounded compartments with electron-transparent matrix (Fig. 2H–L) adjacent to the large posterior vacuole (Fig. 2H, I, K). The 3D reconstruction revealed that each of these compartments contained one crystal of variable size (SI Appendix Movie 5). Less frequently, crystals were found inside the posterior vacuole (Fig. 2I) or in compartments localized near the anterior flagellar pocket (SI Appendix Movie 6), while they were absent from the cytoplasm and other organelles. The crystals had a shape of rhombic prisms (Fig. 2J, M) or asymmetric tabular prismatic structures with pyramidal and pedial terminations (Fig. 2K, N). Although in *L. lanifica* the celestite crystals were mostly lost from the TEM sections, the position of holes and ruptures within them and the analysis by Raman microscopy showed similar localization and size of the crystals to those of *Lacrimia* sp. YPF1808 (Fig. 2D, E, H, I). Likewise, the membrane-bounded compartments were positioned around the posterior vacuole (Fig. 2F), with small asymmetric flattened crystals preserved only occasionally in TEM sections (Fig. 2G).

**Figure 2:**
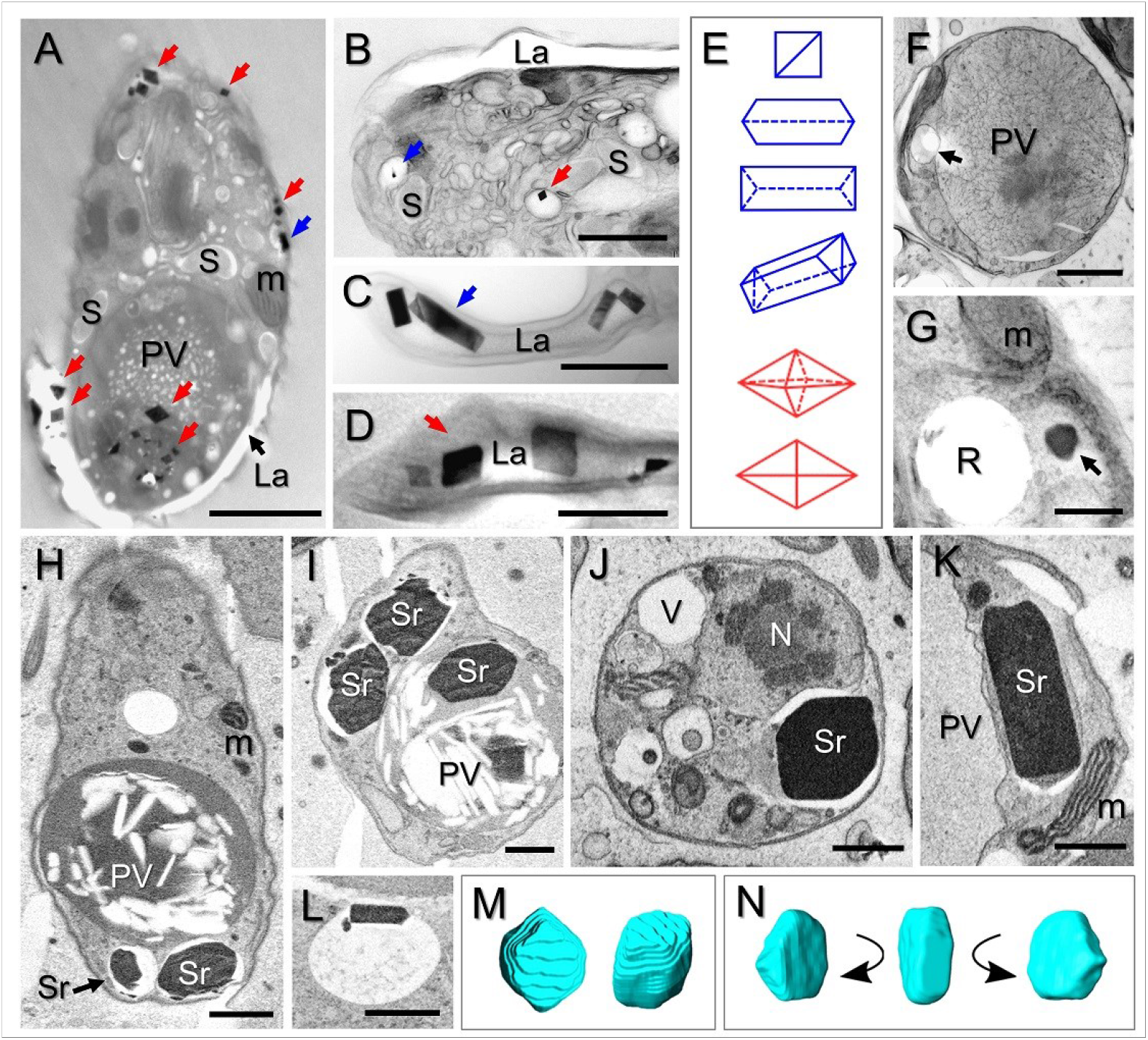
Crystal structures of naturally occurring celestite with possible barite admixtures in diplonemids. TEM of semi-thin sections of *N. karyoxenos* (*A–D*), longitudinally sectioned cell showing bipyramidal (red arrows) and prismatic (blue arrow) crystals inside peripheral lacunae and posterior vacuole (*A*); crystals contained within small vacuoles (*B*); prismatic (*C*) and bipyramidal (*D*) crystals inside lacunae; schematic representation of prismatic (blue) and bipyramidal (red) crystals (*E*). TEM of semi-thin sections of *L. lanifica* JW1601 (*F, G*); cell cross-sectioned through posterior vacuole, *arrow* points to a small membrane-bounded compartment with the hole left after dropped-out crystal (*F*); crystal inside membrane-bounded compartment (*arrow*) and rupture introduced by crystal during sectioning (*G*). SBF-SEM images of *Lacrimia* sp. YPF1808 show celestite crystals inside membrane-bounded compartments (*H–L*); and the large posterior vacuole (*I*). 3D reconstructions of celestite (*M, N*) corresponding to *J* and *K*, respectively, in shape of rhombic prism (*J, M*) and asymmetric tabular prismatic crystal with pyramidal and pedial terminations (*K, N*). PV – posterior vacuole, m – mitochondrion, S – endosymbiotic bacteria, La – lacuna lumen, R – rupture, Sr – celestite crystal, N – nucleus. Scale bar = 2 μm (*A, F*), 1 μm (*B, H–K*) and 500 nm (*C, D, G*).

The presence of celestite crystals (Figs. 1, 2) was further confirmed by elemental analysis using energy-dispersive X-ray spectroscopy in the cryo-SEM-EDX mode of freeze-fractured *Lacrimia* sp. YPF1808 (Fig. 3A, C; SI Appendix Fig. S3) and *N. karyoxenos* cells (Fig. 3B, C; SI Appendix Fig. S4), and by TEM-EDX of whole air-dried cells of *L. lanifica* (Fig. 3E). Atomic percentages estimated by cryo-SEM-EDX analysis were 7.2 % Sr and 7.2 % sulfur (S) as compared to 1.1 % Sr and 1.8 % S in *Lacrimia* sp. YPF1808 and *N. karyoxenos*, respectively. The dominance of C, N, and O atoms can be explained by the presence of ice and signals from other cellular contents obtained from deeper and/or surrounding areas.

**Figure 3:**
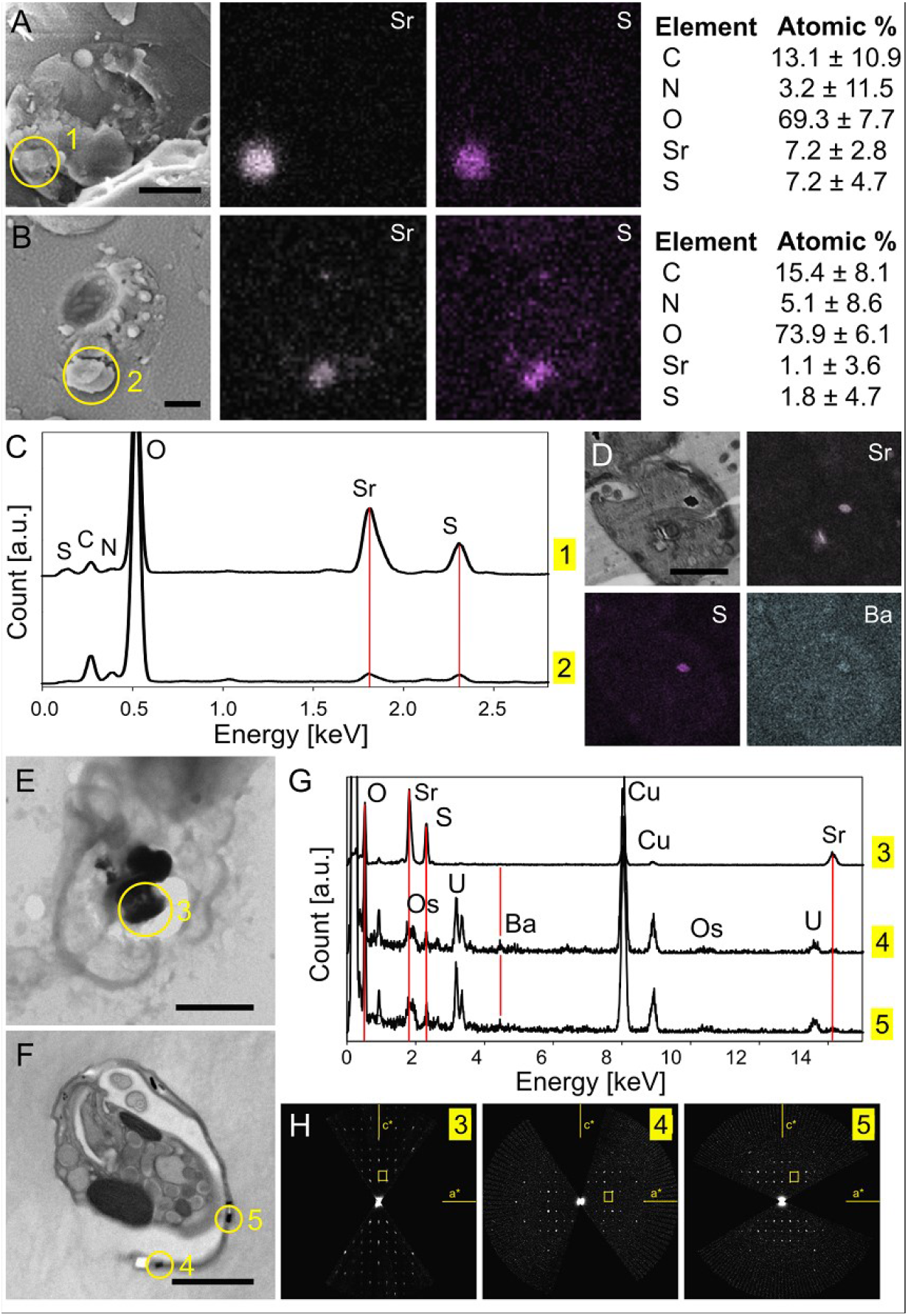
Elemental analysis of diplonemids. Cryo-SEM-EDX images of (*A) Lacrimia* sp. YPF1808 and (*B*) *N. karyoxenos*, complemented with (*C*) EDX elemental spectral analysis obtained from the area marked by circle 1 and 2, respectively. SEM-EDX images of (*D) Lacrimia* sp. YPF1808 showing the presence of S, Sr, and Ba. TEM analysis of celestite microcrystals: (*E*) micrograph of a dried cell of *L. lanifica*, (*F*) micrograph of a semi-thin section of *N. karyoxenos*, (*G*) EDX spectra from the corresponding areas marked 3–5 with red lines highlighting the positions of O, Sr, Ba and S; Cu and C originated from the support grid, Os and U originated from staining compounds, (*H*) electron diffraction: h0l oriented sections through the 3D ED datasets from the corresponding areas shown in (*E*) and (*F*) – celestite unit cell is displayed as a yellow rectangle; scale bar 2 μm.

The identity of celestite crystals in *Lacrimia* sp. YPF1808 (1 μm-thick sections from resin blocks used for SBF-SEM) and *N. karyoxenos* (250 nm-thick resin sections examined by TEM) has been confirmed by SEM-EDX and TEM-EDX, respectively (Fig. 3D, F, G; SI Appendix Fig. S5). Additionally, a significant amount of Ba was detected in the crystals from *N. karyoxenos*. Crystallographic analysis by electron diffraction showed that the diffraction of measured crystals corresponded to celestite structure (isostructural with BaSO_4_) with space group Pnma and lattice parameters *a* = 8.3 Å, *b* = 5.3 Å, *c* = 6.8 Å in *L. lanifica*. Larger lattice parameters (*a* = 8.7 Å, *b* = 5.5 Å, *c* = 7.1 Å) were observed in *N. karyoxenos*, which may be explained by the substitution of Sr^2+^ with larger Ba^2+^ in the structure of celestite.

### Quantitative analysis by ICP-MS and SBF-SEM

SBF-SEM-based 3D reconstructions of *Lacrimia* sp. YPF1808 (SI Appendix Movie 7) showed the presence of celestite crystals in all 20 analyzed cells, ranging from 2 to 16 celestite particles per cell (Fig. 4D). In total, more than 100 crystals were analyzed, with a volume ranging from 0.017 to 7 μm^3^ (Fig. 4C). The impact of the measured celestite contents on the overall cell density ranged from 0.05 to 9 %, with an average of 1.3±0.5 % (Fig. 4E). The calculations were based on the measured volumes, known density of celestite (3.9 g·cm^−3^) and common cellular densities of 0.985–1.156 g·cm^−3^ reported elsewhere [48].

**Figure 4:**
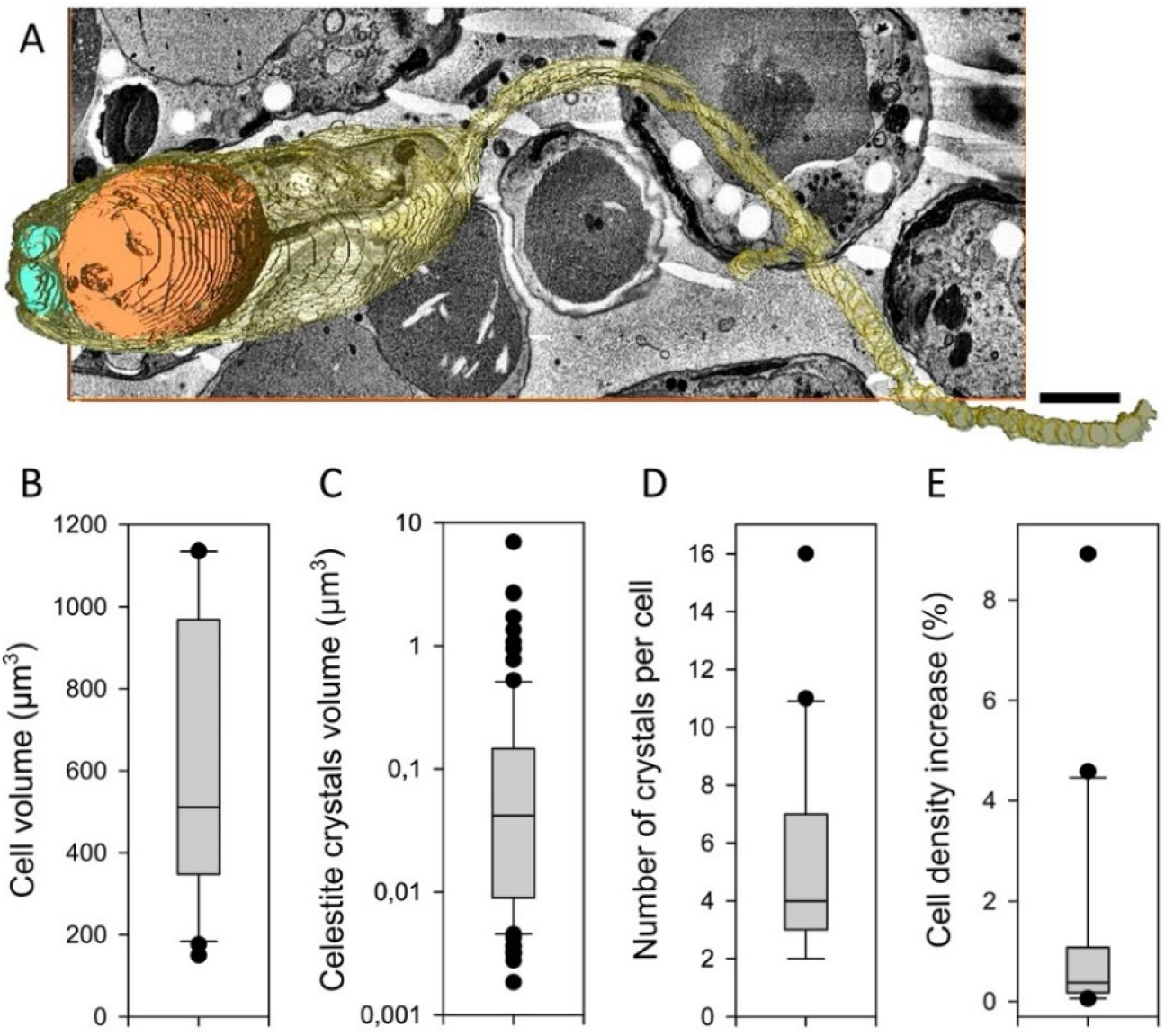
SBF-SEM imaging of celestite crystals in *Lacrimia* sp. YPF1808. (*A*) 3D reconstruction of a cell: cytoplasm in yellow, large posterior vacuole in orange, and celestine crystals in cyan; scale bar = 1 μm. Descriptive analysis of measured data, cells N=20, crystals N=106: (*B*) distribution of cell volume, (*C*) distribution of celestite crystals volume on a log scale, (*D*) number of celestite crystals per cell, (*E*) impact of celestite crystals on cell density.

The lack of celestite crystals in other analyzed species (*Diplonema japonicum, Paradiplonema papillatum* and *Rhynchopus* sp. YZ270) was consistent with minute ^88^Sr content measured by ICP-MS. The high values of ^88^Sr in *N. karyoxenos, Lacrimia* sp. YPF1808 and *L. lanifica* corresponded to the abundance of intracellular crystals detected by Raman microscopy, TEM and SBF-SEM. Since direct measurement of the dry mass was impossible due to the inevitable presence of salts from the medium, the elemental composition analysis by ICP-MS was calculated in atoms·cell^−1^, or fmol·cell^−1^. To calculate Sr and Ba content per dry mass, the latter was subsequently estimated by quantitative phase imaging using holographic microscopy (Table 1). The ^88^Sr amounts ranged from 0.01 fmol·cell^−1^ in *P. papillatum* to 5500±570 fmol·cell^−1^ in *N. karyoxenos*, corresponding to 340±38 mg·g^−1^. *Lacrimia* sp. YPF1808 and *L. lanifica* are also potent ^88^Sr accumulators with 370±58 fmol·cell^−1^ (130±25 mg·g^−1^) and 54±8 fmol·cell^−1^ (64±13 mg·g^−1^), respectively. Depending on the species, the intracellular concentration of ^88^Sr was 1,200 to almost 10,000 times higher than in the surrounding medium (Table 1).

**Table 1:** ICP-MS quantification of ^88^Sr and Ba in diplonemids. The dry weight was measured by quantitative phase imaging (n = 150 cells). Results of ICP-MS are displayed as mean values of biological triplicates with standard error of the mean. All figures are rounded to two significant numbers. The amounts of ^88^Sr and Ba per dry mass (mg·g^−1^) was logarithmically transformed and statistically analyzed by one-way ANOVA *p* < 0.001, with Tukey’s post-hoc test significant differences on the level *p* < 0.05 displayed in column *p*.

| Species | *D. japonicum* | *D. papillatum* | *Rhynchopus*<br>sp. YZ270 | *L. lanifica*<br>JW1601 | *Lacrimia* sp.<br>YPF1808 | *N. karyoxenos* |
| --- | --- | --- | --- | --- | --- | --- |
| Dry weight of cell (pg) | 109 ± 2.9 | 68.0 ± 2.0 | 74.2 ± 2.7 | 74.1 ± 3.5 | 248.1 ± 8.2 | 1,421 ± 11 |
| Sr (fmol·cell<sup>-1</sup>) | 0.22 ± 0.01 | 0.013 ± 0.001 | 0.08 ± 0.02 | 54.4 ± 8.4 | 366 ± 58 | 5,530 ± 570 |
| Sr (mg·g<sup>-1</sup>) | 0.17 ± 0.10 | 0.02 ± 0.01 | 0.10 ± 0.07 | 64 ± 13 | 129 ± 25 | 340 ± 38 |
| $p$ | **a** | **b** | **a** | **c** | **d** | **e** |
| SrSO<sub>4</sub> (mg·g<sup>-1</sup>) | n/a | n/a | n/a | 135 | 271 | 715 |
| Concentration of Sr (folds)* | n/a | n/a | n/a | 1,520 ± 680 | 1,240 ± 550 | 10,000 ± 3,300 |
| Ba (fmol·cell<sup>-1</sup>) | n/a | n/a | n/a | 2.0 ± 0.5 | 65 ± 10 | 1,200 ± 130 |
| Ba (mg·g<sup>-1</sup>) | n/a | n/a | n/a | 3.7 ± 1.1 | 35.8 ± 6.8 | 116 ± 14 |
| $p$ | – | – | – | **a** | **b** | **c** |
| BaSO<sub>4</sub> (mg·g<sup>-1</sup>) | n/a | n/a | n/a | 6.3 | 61 | 198 |
| Concentration of Ba (folds)* | n/a | n/a | n/a | 1,130 ± 660 | 4,300 ± 1,900 | 42,000 ± 14,000 |
| Total Ba,Sr(SO<sub>4</sub>) (mg·g<sup>-1</sup>) | **n/a** | **n/a** | **n/a** | **141** | **332** | **913** |

n/a – not analyzed
\* – calculated as atoms·cell<sup>-1</sup> based on analyzed cell pellets relative to the theoretical value in atoms·cell<sup>-1</sup> originating from the culture medium alone (8 mg·l<sup>-1</sup> Sr<sup>2+</sup> and 0.64 mg·l<sup>-1</sup> Ba<sup>2+</sup>); in *N. karyoxenos*, the amount of accumulated Ba exceeds total Ba available per volume of culture medium due to repeated passages of pelleted cells into fresh medium

Compared to the massive accumulation of ^88^Sr, the naturally co-occurring Ba was present in much lower amounts (Table 1), slightly above the detection limit of TEM-EDX analysis (Fig. 3G), possibly reflecting 12.5 times lower Ba^2+^ concentration in the seawater growth medium (SI Appendix Table S1). Nevertheless, the cells concentrated Ba^2+^ on average 1,000 to over 42,000 times above the level in the growth medium (Table 1), reaching 1,200±130 fmol·cell^−1^ (120±14 mg·g^−1^) in *N. karyoxenos*, with lower values in *Lacrimia* sp. YPF1808 (65±10 fmol·cell^−1^ or 36±7 mg·g^−1^) and *L. lanifica* (2±1 fmol·cell^−1^ or 4±1 mg·g^−1^). Altogether, (Ba,Sr)SO_4_ accumulation reached 91 %, 33 % and 14 % of dry weight in *N. karyoxenos, Lacrimia* sp. YPF1808 and *L. lanifica*, respectively. All strains showed significantly different levels of accumulated Sr or Ba contents (Table 1).

### Ba^2+^ loading experiments and elimination of Sr^2+^ and Ba^2+^ from the medium

Pure barite has not been detected by Raman microscopy in any of the examined species, apparently because in the growth medium, Ba^2+^ is ~12.5 times less abundant than Sr^2+^ (SI Appendix Table S1). However, upon cultivation in artificial seawater loaded with equimolar amounts of Ba^2+^ and Sr^2+^, we observed the occurrence of barite in all three diplonemids (Fig. 5). This experiment showed biocrystallization of all mineral combinations: pure barite (Raman marker at 988 cm^−1^) and celestite (1,000 cm^−1^), as well as mixed forms of (Ba,Sr)SO_4_ (991 cm^−1^; strontiobarite or baritocelestite) (Fig. 5), revealing that diplonemids do not show a strong preference towards the accumulation of either element.

**Figure 5:**
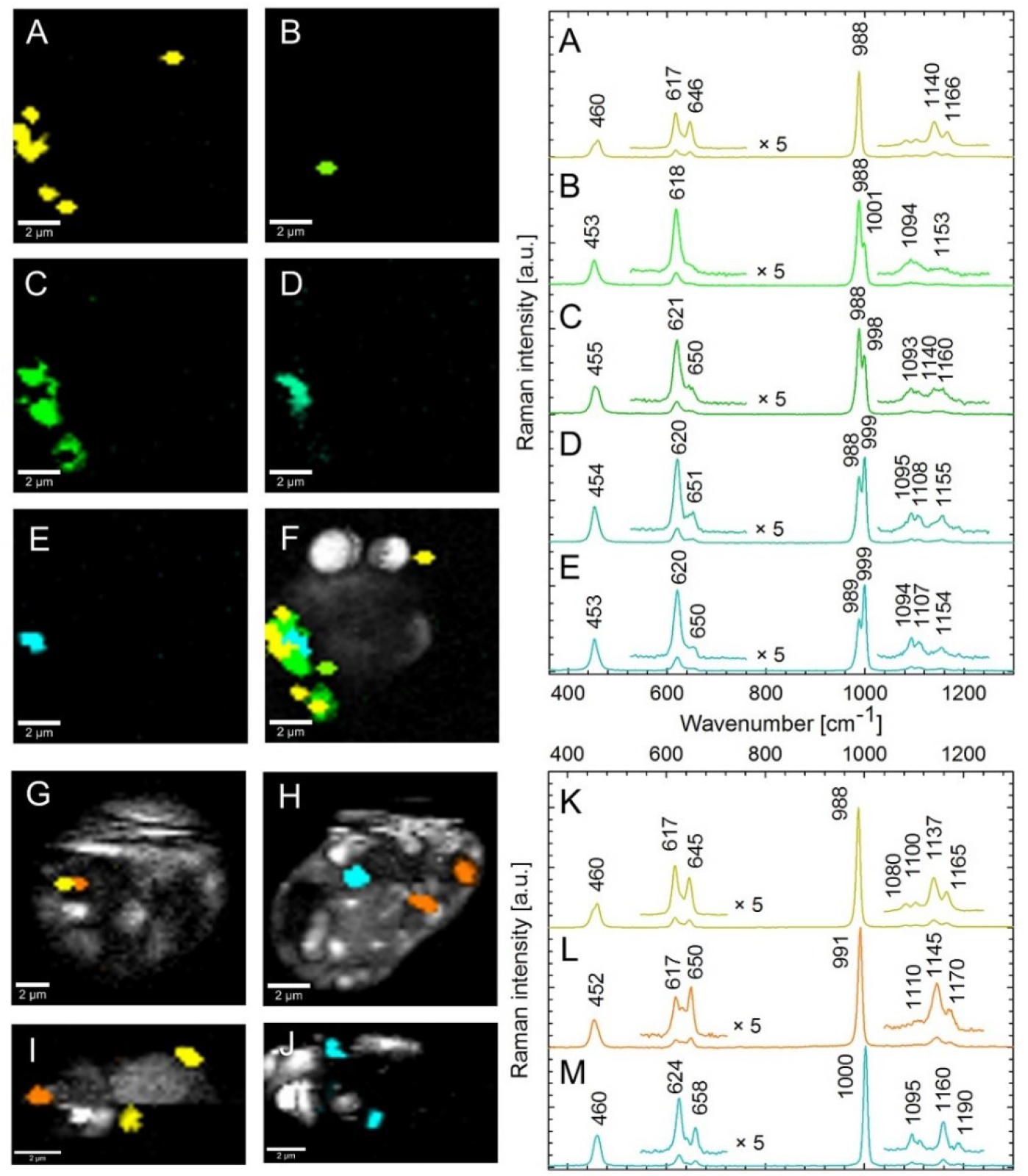
Raman maps and spectra of barite and celestite crystals in diplonemids cultivated with equimolar amounts of Sr^2+^ and Ba^2+^ in the medium. A single cell of *N. karyoxenos (A–F*) containing co-crystallized fractions dominated by celestite in the central part (*E* in cyan) with gradually overlapping barite (*B–D* in shades of green) towards the periphery of its pure fraction (*A* in yellow), merged Raman map of *A–E (F*). The limited spatial resolution of Raman microscopy does not allow distinguishing conglomerate of pure-species microcrystals from a single crystal with a variable elemental composition. Two cells of *Lacrimia* sp. YPF1808 (*G, H*) and two cells *L. lanifica (I, J*) contain pure barite (*K* in yellow) and celestite (*M* in blue) and homogenously mixed crystals of (Ba,Sr)SO_4_ (*L* in orange).

Both *L. lafinica* and *Lacrimia* sp. YPF1808 contained mostly mixed crystals of (Ba,Sr)SO_4_. In *N. karyoxenos*, structured co-crystallization of the two pure components occurred, in which celestite prevailed in the central part of crystals (999 cm^−1^), while barite dominated at their periphery (988 cm^−1^), gradually overlapping each other (Fig. 5C–F). Additionally, after several passages in the artificial medium without Ba^2+^ and Sr^2+^ (SI Appendix Table S1), barite and celestite were no longer detectable by Raman microscopy. The lack of Ba^2+^ and Sr^2+^ did not result in altered morphology or growth impairment. Moreover, in the absence of Sr^2+^ and Ba^2+^, we did not observe any Ca^2+^/Mg^2+^-containing crystals despite high concentration of both elements in the medium.

### Feeding experiments

Fecal pellets are an important agent mediating sedimentation of biogenically-accumulated minerals to the sea floor: large aggregates of crystals held together by undigested fecal organic matter enable their fast sinking, thus preventing from dissolution of micrometer-sized crystals in the water column, which is undersaturated for barite and celestite [11, 23]. To experimentally address whether zooplankton feeds on diplonemids and whether their barite and celestite crystals are carried into fecal pellets, we incubated *N. karyoxenos* and *Lacrimia* sp. YPF1808 with freshly captured filter-feeding marine copepods *Centropages typicus*, *Temora longicornis* and *Acartia* sp., starved for 12 h prior to the experiment. After 5-day co-cultivation, we determined by Raman microscopy that the fecal pellets contained copious amounts of celestite derived from diplonemids (Fig. 6B). In the control system of the same population of copepods fed with freshly collected marine plankton, the fecal pellets contained undigested chlorophyll, carotenoids with remnants of lipids, calcite particles, and contaminating polystyrene microplastic particles, but lacked barite and celestite crystals (Fig. 6B).

**Figure 6:**
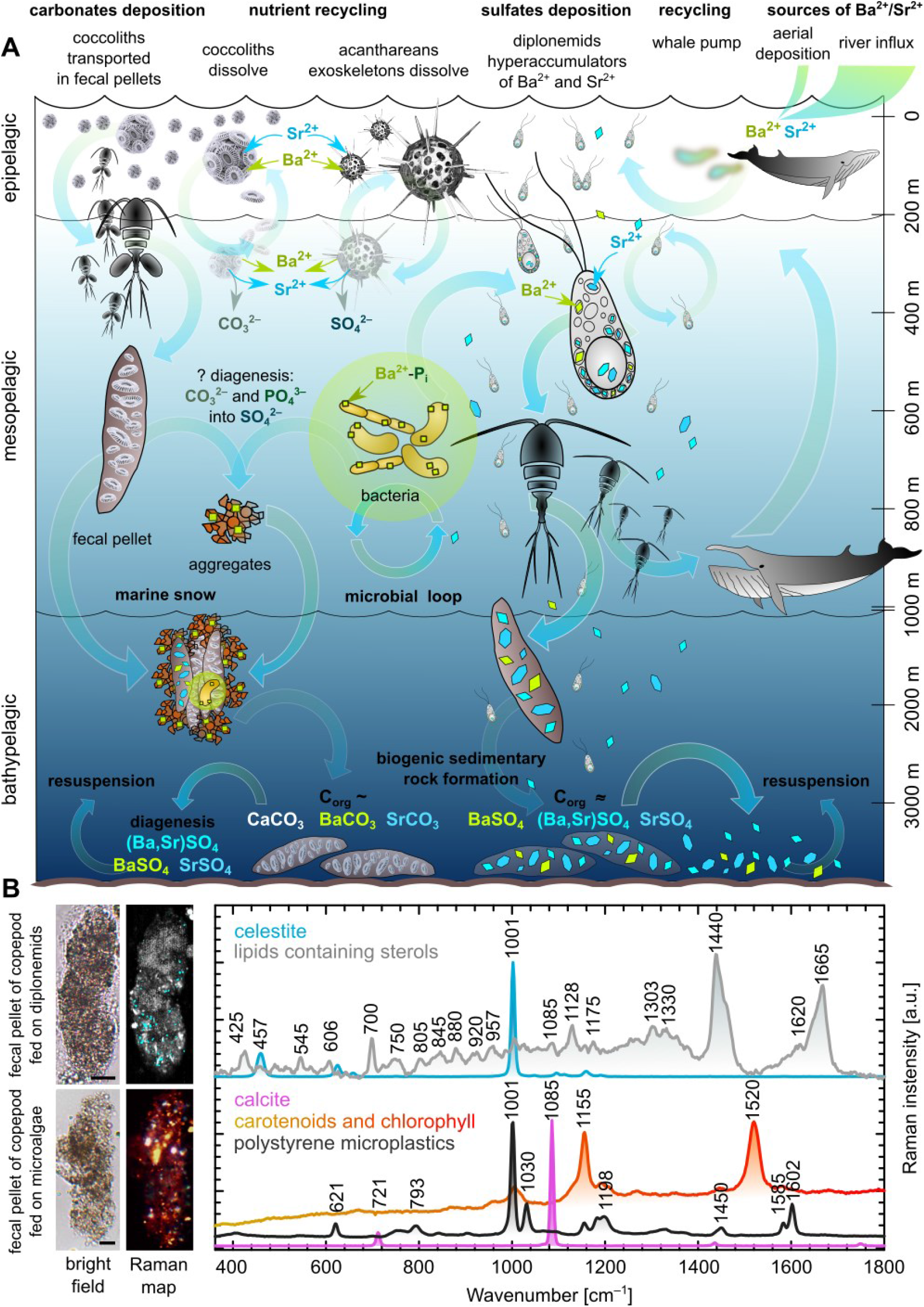
Schematic representation of the biological influence of Ba^2+^ and Sr^2+^ cycling in the oceans. (*A*) Hypothetical scenario of trace elements inlet, plankton uptake, recycling, and sedimentary deposition. The major source of trace elements is driven by river influx and less prominently by aerial deposition [9] and is mostly balanced by the same amount of total deposition in sediments [23]. A great proportion of these trace elements is being recycled by living organisms after release from dead cells. General nutrient flux goes from the surface primary producers to the zooplankton scattered throughout the water column to the deep sea and is bypassed by the “whale pump” bringing the nutrients from the depth to the surface where whales defecate [59]. Acantharea (image adapted from [69]) uptake a substantial part of Sr^2+^ and Ba^2+^, which is further recycled in upper 400 m [24]. Coccolithophorids build their scales from calcium carbonate with minor amounts of Ba^2+^ and Sr^2+^ that are proportional to the seawater contents [19], being partially recycled upon dissolution or transported to the marine sediments in fecal pellets [23]. The Ba^2+^ accumulated by bacteria sediments in the aggregates of marine snow [25]. We highlight diplonemids as potential players in the marine cycle of both elements and drivers of biogenic formation of celestite and barite crystals found in suspended matter everywhere in the world ocean [23]. (*B*) Raman microscopy analysis of fecal pellets produced by copepods experimentally fed with diplonemids, which contain celestite crystals (cyan) and undigested lipids including sterols (grey). In the control samples of copepods fed by microalgae, fecal pellets contained undigested carotenoids, chlorophyll (yellow-orange-red) and calcite particles (pink) with unexpected particles of polystyrene microplastics (measured as a single spectrum – not shown on the Raman map). Scale bars 20 μm.

## DISCUSSION

The most studied biominerals in protists are extracellular calcite scales of haptophytes and silicate frustules of diatoms, while studies on intracellular mineral crystals are far less common [53]. After more than a century since the skeletons of marine acanthareans and freshwater streptophytes were found to contain celestite [12] and barite [54], respectively, we have identified potent accumulators of Ba^2+^ and Sr^2+^ in an unexpected group of eukaryotes, the diplonemids. These planktonic heterotrophic flagellates are highly abundant in the nutrient-rich mesopelagic waters, which are important components of the biological pump of the carbon cycle [32, 33, 55]. The high capacity of intracellular Sr^2+^ and Ba^2+^ accumulation in diplonemids outperforms that of any other reported organisms [10, 15, 18, 56–58]. Indeed, while the intracellular concentration of Sr^2+^ in the most efficient accumulators known so far (yeasts, desmids and cyanobacteria) reaches a maximum of 220 mg·g^−1^ per dry weight [1, 10, 57], *N. karyoxenos* contains as much as 340 mg·g^−1^ Sr^2+^ together with 120 mg·g^−1^ Ba^2+^, which in the form of sulfate represents 90 % of the cellular dry mass, pointing to the unique Ba^2+^ and Sr^2+^ accumulation capacity of this diplonemid, while both *Lacrimia* species are only slightly less potent in this respect (Table 1). It has to be mentioned that the cultivation media differ from the environmental conditions when Ba^2+^ and Sr^2+^ are considered, and the same applies for cell densities that are higher by orders of magnitude than in the water column (SI Appendix Table S1). Still, the capability of the globally most diverse deep-sea pelagic diplonemids [32, 33] to form barite and celestite crystals could not be analyzed due to their unavailability in culture, it is reasonable to anticipate it as an analogy to similar system found in zygnematophytes [14].

The massive formation of barite and celestite in diplonemids documented herein, allows us to propose a novel view of the Ba^2+^/Sr^2+^ cycling in the ocean (Fig. 6). Celestite-forming acanthareans are considered as key players in the upper 400 m of the ocean, yet without contributing to the sedimentary rock formation, as their skeletons dissolve upon decay of their cells [24]. Coccolithophorids and bacteria produce carbonates [58] and/or phosphates [25] of Ba^2+^/Sr^2+^, which can also be converted to sulfates either on the polysaccharides of bacterial cell walls or in the microenvironment of decaying matter of marine-snow aggregates in the process of diagenesis [25]. The majority of biogenic particulate barite and celestite is being recycled by simple dissolution [24], microbial loop [25], resuspension of sediments [23] or whale pump recycling [59]. However, the overall influx into the system is balanced by the sedimentary deposition [9, 23] that might have a biological driver. Seminal work of Dehairs et al. [23] scrutinized all potential sources of particulate barite from precipitation on decaying organic matter in sulfate-enriched microenvironments to Ba^2+^ incorporation in siliceous plankton that got disproved by series of laboratory experiments, ultimately favored the biogenic origin of particulate barite from as-yet-unknown microorganisms. The particulate barite maxima in high-productivity areas below the euphotic zones led to the prediction of active precipitation by planktonic heterotrophs that remained unknown for more than four decades. These predictions [23] nicely correlate with our measurement in diplonemids: micron-sized barite and celestite crystals of variable Ba-Sr ratio (Fig. 2–5) are scattered throughout the water column of world’s ocean, with the highest prevalence in the mesopelagic zone [32]. Moreover, such particulate barite is often found in fecal pellets and aggregates of marine snow, and finally, in the sediments [23, 27, 32] and here we showed that that celestite occurring in diplonemids can be transferred to fecal pellets of filter-feeding copepods (Fig. 6).

Despite their abundance and extreme diversity, diplonemid flagellates remain a poorly known group of protists [34], that are abundant from the surface to the deep sea, with a wide peak in the mesopelagic and bathypelagic zones [32, 33, 60]. Only over a dozen species of these naked fragile heterotrophs can be cultivated in the laboratory, with most of them being included in this study. Our findings suggest that diplonemids are direct producers of pelagic barite and celestite. In general, the most effective transport mechanism of biominerals to the sea floor is in the fecal pellets of zooplankton that consumed planktonic microbes possessing indigestible barite and/or celestite crystals [23]. We experimentally showed that barite and celestite, derived from diplonemids, is indeed carried to fecal pellets of copepods (Fig. 6). Thus, diplonemids may be involved in Ba^2+^/Sr^2+^ cycling and/or in sedimentary deposition of barite and celestite. Since these protists likely emerged already during the Neoproterozoic era (590–900 MYA) overlapping with the Ediacaran period [61], their impact on biogenic marine sediments may cover several geological eras. The coccolithophores appeared around the same time as diplonemids, yet the onset of carbonate biomineralization was timed to ~200 MYA [62].

Interestingly, when both trace elements are provided in equimolar concentrations, diplonemids form pure celestite and barite and/or mixed forms of (Ba,Sr)SO_4_, apparently not discriminating one element over the other. Hence, we explain the higher content of Sr^2+^ over Ba^2+^ inside the crystals by the higher availability of the former element in seawater. Although the mechanisms behind intracellular accumulation of Sr^2+^ and Ba^2+^ are largely unknown, it has been suggested that mineral crystals typically occur in membrane-bounded compartments or vacuoles, in which they are formed from supersaturated solutions *via* precisely regulated nucleation [15]. The Sr^2+^ uptake and transportation within eukaryotic cells have been shown to occur *via* the commonly present transporters of divalent cations, *i.e*. the Ca^2+^ uniporter and H^+^/Ca^2+^ antiport [63, 64]. The diplonemid nuclear genome is not yet available, but these transporters have been documented in the related kinetoplastid *Trypanosoma brucei* [65]. Although the reported affinity to Ca^2+^ and Sr^2+^ is usually comparable [63, 64], some organisms including diplonemids clearly favor Ba^2+^ and Sr^2+^ over Ca^2+^ [10]. When such vacuoles contain sulfate solutions, they may function as a “sulfate trap” for those cations that precipitate easily in the presence of sulfates [2]. At the same time, we did not observe CaSO_4_ or any of its forms (gypsum, bassanite, anhydride, etc.) even though the concentration of Ca^2+^ in the cultivation medium or in the environment is several orders of magnitude higher than that of Sr^2+^ and Ba^2+^.

Dense celestite and barite of 3.9 g·cm^−3^ and 4.5 g·cm^−3^, respectively, were repeatedly reported as statoliths in ciliates or charophytic algae [13–15]. In comparison to the seawater density of 1.03 g·cm^−3^ and typical cell density range between 0.985 and 1.156 g·cm^−3^, the heavy crystals may help maintain appropriate buoyancy by counterbalancing light lipid droplets (0.86 g·cm^−3^) [48, 66]. Indeed, the impact of celestite crystals is substantial, since they may increase the overall density of *Lacrimia* sp. YPF1808 and *N. karyoxenos* by up to 9 % and 27 %, respectively. According to Stokes’ law for small particles of low Reynolds numbers, the barite-celestite ballasting can significantly increase the sedimentary velocity for up to 50–200 m per month or 0.5 to 2 km per year. Hence, while the function of biomineralization in diplonemids remains unknown, we speculate that they may benefit from gravitropic sensing, which would allow directed movement and/or enable passive sedimentation. Another intriguing impact of barite and celestite is associated with their propensity to strong absorption of UV and blue light [67]. Hence, in surface waters, these minerals may contribute to UV protection. It is reasonable to assume that by forming celestite, protists adjust their inner osmolarity – the principle analogical to formation of other cell inclusions, such as oxalate, calcite or polyphosphate, that are either dissolved and osmotically active or crystallized/polymerized and osmotically inactive inside a vacuole [15, 68]. Based on the ability of some diplonemids to store massive amounts of barite and celestite, we speculate that they may qualify as hitherto unknown yet impactful players of Ba^2+^/Sr^2+^ flow through the food web, eventually influencing the sedimentary records.

## ACKNOWLEDGEMENTS

This work was supported by the Czech Ministry of Education ERD Funds projects OPVVV 16_019/0000759 (to JL), 15_003/0000336 (to HK and BSNH) and 16_013/0001775 (to DT, JT and MV), ERC CZ LL1601 (to JL), the Czech Bioimaging grant LM2018129 (to JP, DT, JT and MV), the Czech Science Foundation grant 21-26115S (to JP and PM), and the Gordon and Betty Moore Foundation GBMF9354 (to JL). We acknowledge CzechNanoLab Research Infrastructure (LM2018110) and the Light Microscopy Core Facility (LM2018129, 18_046/0016045) for help with holographic microscopy.

## Author contributions

DT, JP and JL designed research; DT, JP, JT, MV, BSNH, RS, and MK performed research; JP, DT, JT, MV, HK, PM and JL analyzed data; JT, MV, HK, PM, RS, and MK contributed reagents/analytic tools; JP, DT and JL wrote the paper.

## Declaration of interests

We declare no conflict of interest.

## Data Availability Statement

All data generated or analyzed during this study are included in this published article and its supplementary information files.

## Movie legends

**Movie 1.** *N. karyoxenos* under bright field DIC microscopy showing fast moving celestite crystals in lacunae marked by arrows.

bit.ly/3qT5gsN

**Movie 2.** *Lacrimia* sp. YPF1808 under bright field DIC microscopy showing light-polarizing celestite crystals marked by an arrow.

bit.ly/3rO6gxX

**Movie 3.** *N. karyoxenos* under bright field polarization microscopy showing light-polarizing celestite crystals marked by arrows.

bit.ly/3nNfitN

**Movie 4.** *Lacrimia* sp. YPF1808 under bright field polarization microscopy showing light-polarizing celestite crystal marked by an arrow.

bit.ly/3Iwe2Df

**Movie 5.** Visualization of serial sections through the celestite-containing cell of *Lacrimia* sp. YPF1808 taken by SBF-SEM, celestite crystals marked by arrows. PV – posterior vacuole.

bit.ly/3fT0wxi

**Movie 6.** Visualization of serial sections through the celestite-containing cell of *Lacrimia* sp. YPF1808 taken by SBF-SEM, celestite crystals marked by arrows. N – nucleus, FP – flagellar pocket.

bit.ly/3qUBi81

**Movie 7.** 3D reconstruction of a single cell *via* SBF-SEM: cytoplasm in yellow, large posterior vacuole in orange and celestine crystals in cyan.

bit.ly/3FU8X64

**All the supplementary movies are deposited here:** 10.6084/m9.figshare.20154200

## Supplementary Information

**Fig. S1.**
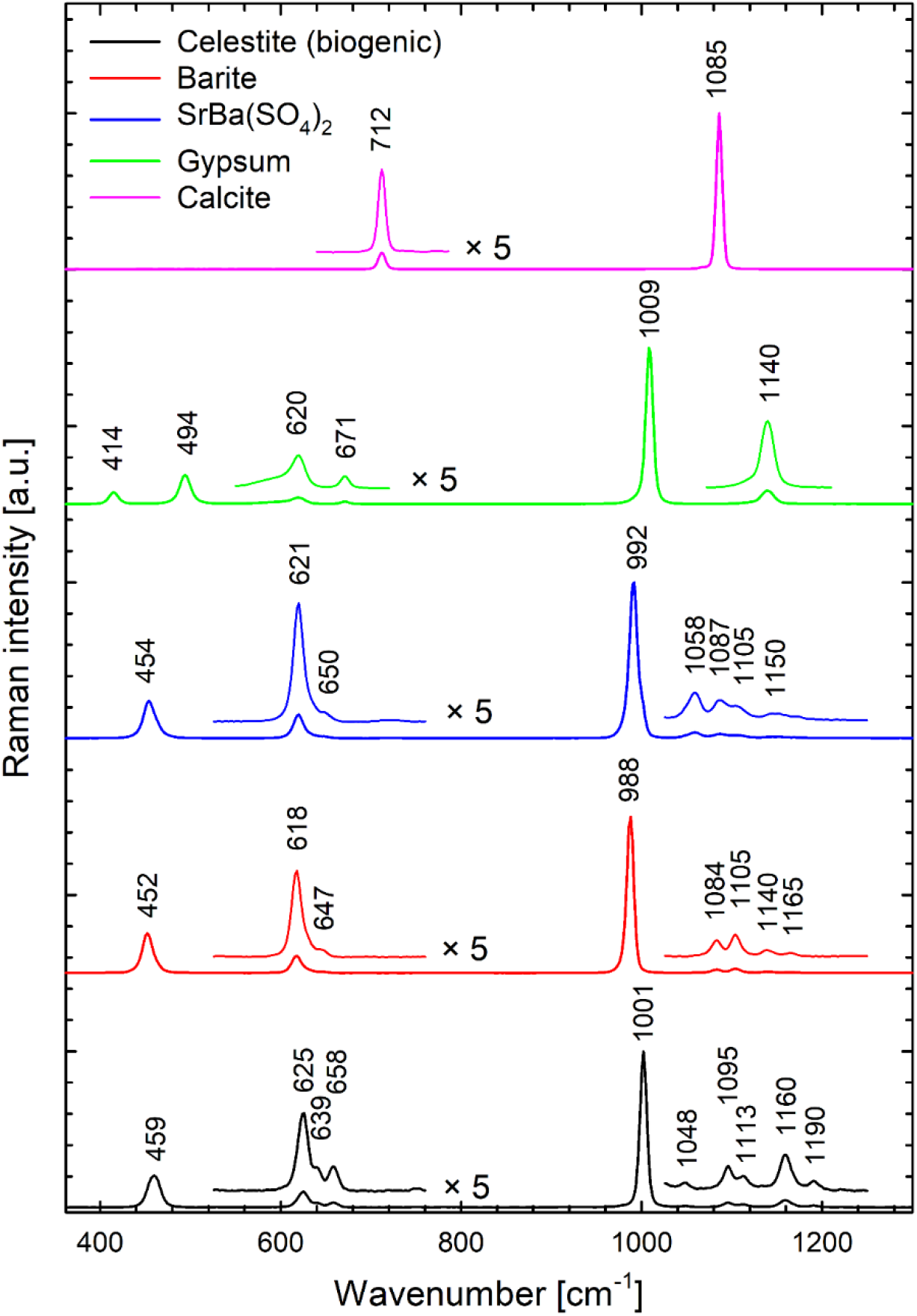
Typical Raman spectrum of biogenic celestite crystals compared with the spectra of chemically prepared barite, equimolar precipitate of SrSO_4_ and BaSO_4_, gypsum (CaSO_4_) and calcite (CaCO_3_). Each complex of SO_4_^2-^ with Sr^2+^, Ba^2+^ or Ca^2+^ can be unambiguously identified by the position of the most intense Raman band at around 1000 cm^−1^ belonging to the symmetric *v*_1_ vibrational mode of oxygen atoms within SO_4_^2-^ tetrahedron.

**Fig. S2.**
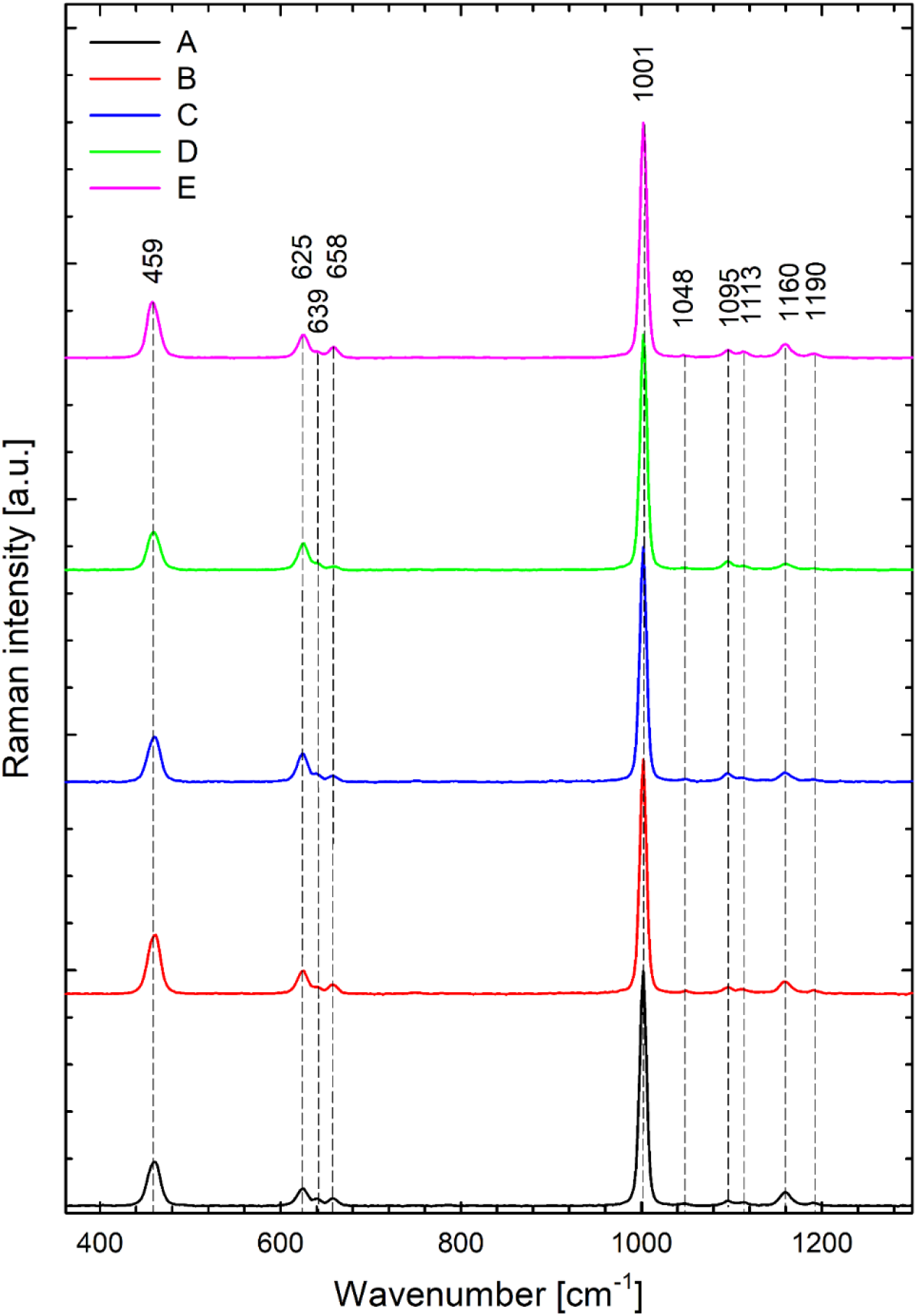
Typical Raman spectra (A–E) of biogenic celestite crystals found in various *Diplonema* cells. As concerns Raman frequencies, the spectra are virtually identical, however minor spectral variability in the relative intensities of Raman bands can be observed. Similar variability was observed also for mineral celestite and for chemically prepared SrSO_4_ precipitates.

**Fig. S3.**
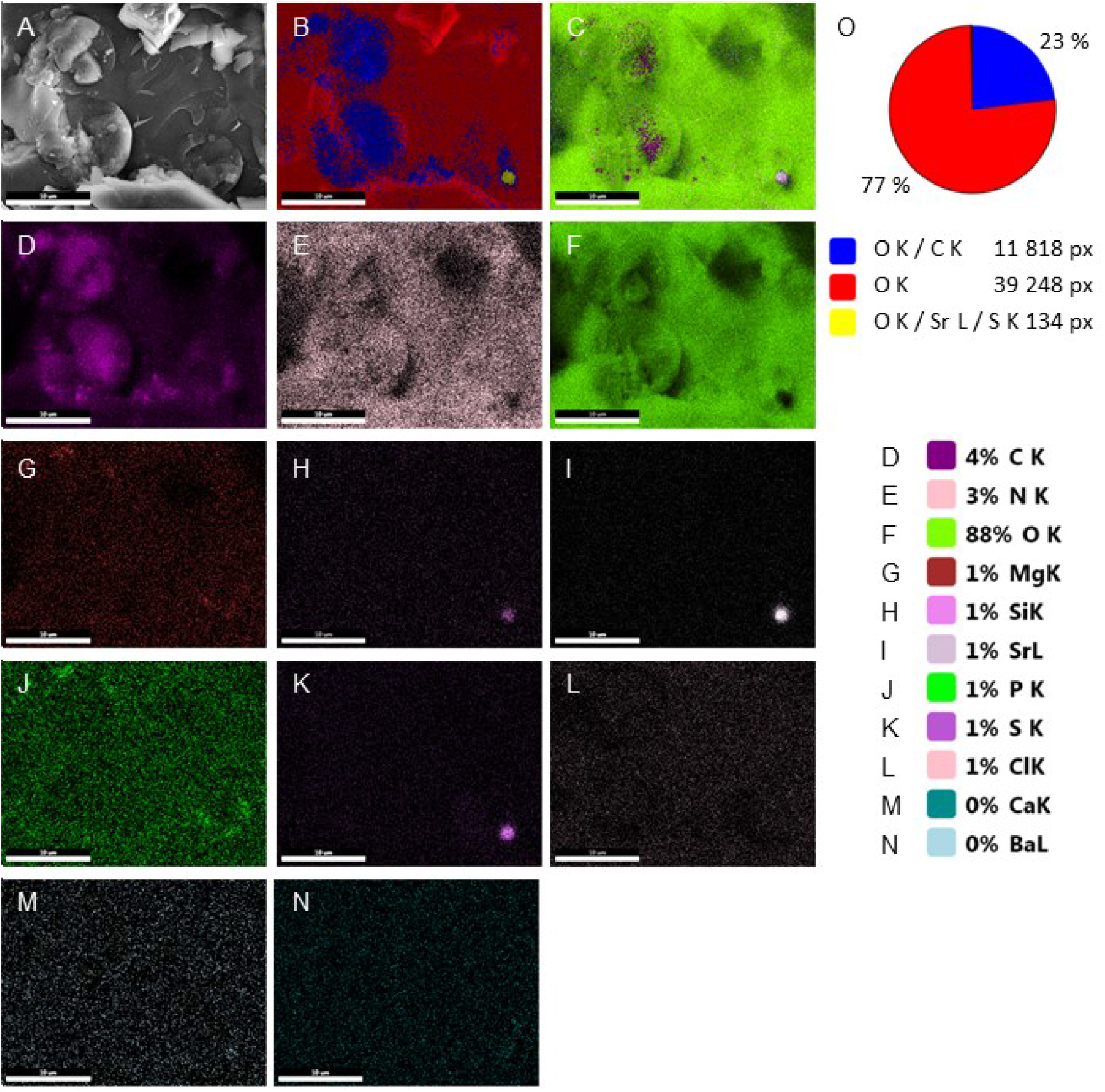
Cryo-SEM-EDX analysis of *Lacrimia* sp. YPF1808 showing the elemental analysis of freeze-fractured samples, A – SEM micrograph, B – merged image of the most common elements, C – merged image of all elements present, D – map of carbon K line, E – map of nitrogen K line, F – map of oxygen K line, G – map of magnesium K line, H – map of silicon K line, I – map of strontium L line, J – map of phosphorus K line, K – map of sulfur K line, L – map of chlorine K lines, M – map of calcium K line, N – map of barium L line, O – pie chart of the most common elements detected in the area with particular pixel counts, scale bar 10 μm.

**Fig. S4.**
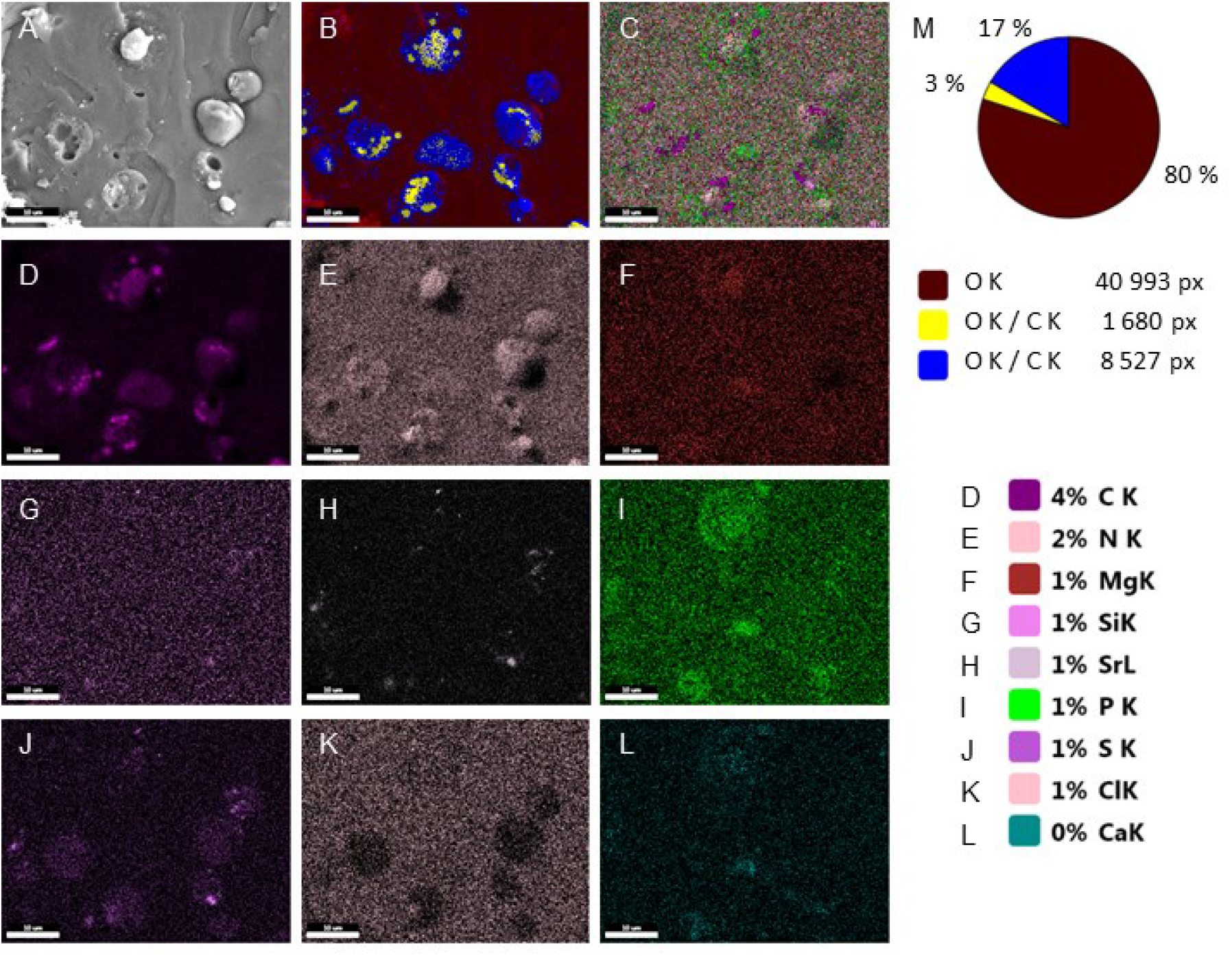
Cryo-SEM-EDX analysis of *N. karyoxenos* showing the elemental analysis of freeze-fractured samples, A – SEM micrograph, B – merged image of the most common elements, C – merged image of all elements present, D – map of carbon K line, E – map of nitrogen K line, F – map of magnesium K line, G – map of silicon K line, H – map of strontium L line, I – map of phosphorus K line, J – map of sulfur K line, K – map of chlorine K lines, L – map of calcium K line, M – pie chart of the most common elements detected in the area with particular pixel counts, scale bar 10 μm.

**Fig. S5.**
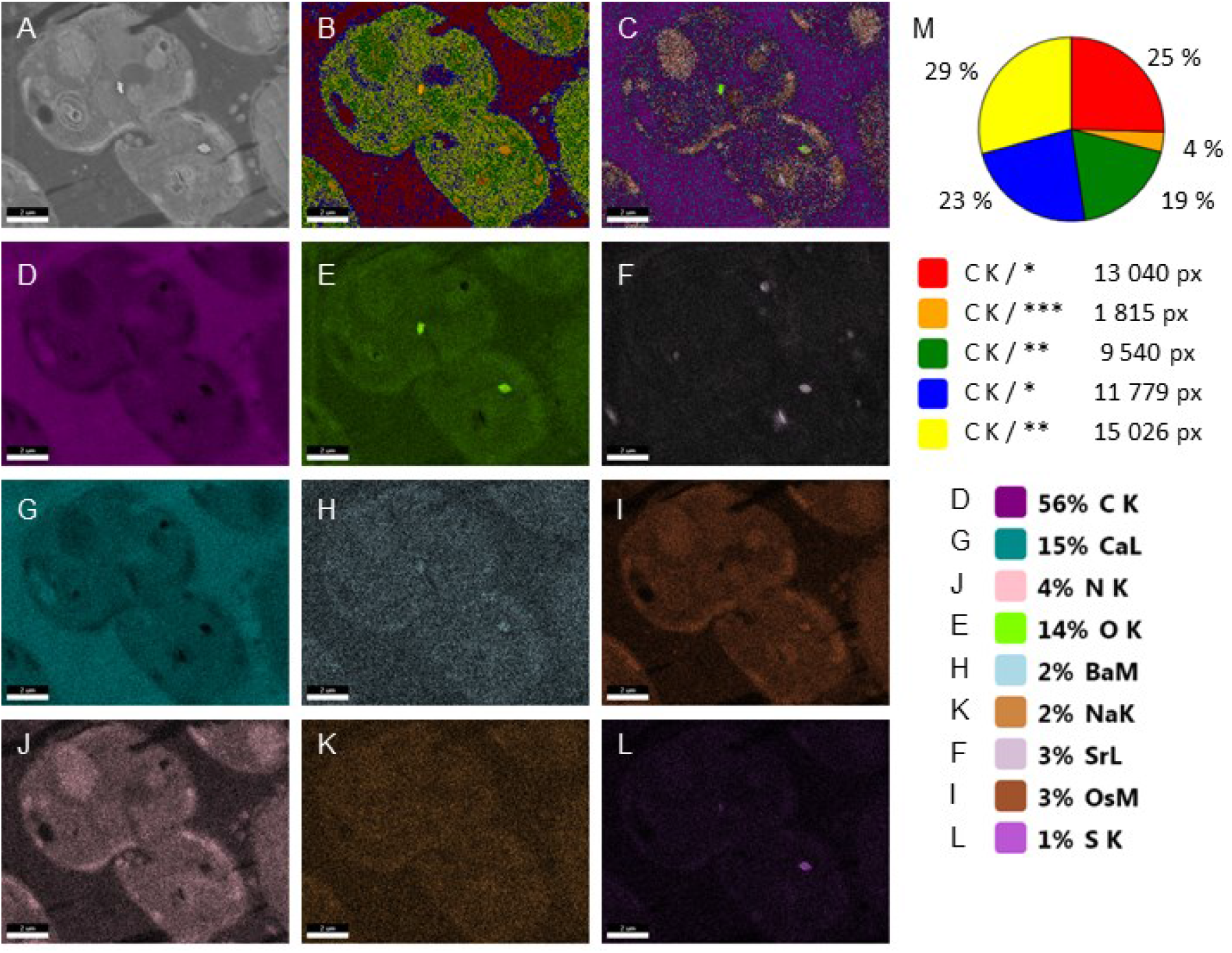
SEM-EDX analysis of *Lacrimia* sp. YPF1808 showing the elemental composition of samples used for 3D reconstruction by SBF-SEM, A – electron-micrograph, B – merged image of the most common elements, C – merged image of all elements present, D – map of carbon K line, E – map of oxygen K line, F – map of strontium L line, G – map of calcium L line, H – map of barium M line, I – map of osmium M line, J – map of nitrogen K line, K – map of sodium K line, L – map of sulfur K lines, M – pie chart of the most common elements detected in the area with particular pixel counts, * represents O K / Ca L, ** represents O K/ Ca L / N K, *** represents O K / Ca L / N K / Sr L / Os M, scale bar 2 μm.

**Table S1.** ICP-MS analysis of seawater growth medium and artificial medium lacking Ba^2+^ and Sr^2+^ sources. Mean of three technical replicates (±SD of Mean).

| Element | unit | Standard seawater growth medium |  | Artificial seawater medium lacking sulfates, Ba and Sr * |  |
| --- | --- | --- | --- | --- | --- |
|  |  | mean | SD | mean | SD |
| Mg | ppb | 1 710 000 | 69 000 | 3 190 000 | 120 000 |
|  | nM | 71 200 000 | 2 900 000 | 132 900 000 | 4 900 000 |
| K | ppb | 1 360 000 | 190 000 | 1 291 000 | 52 000 |
|  | nM | 34 800 000 | 4 900 000 | 33 100 000 | 1 300 000 |
| Ca | ppb | 513 000 | 34 000 | 545 000 | 18 000 |
|  | nM | 11 650 000 | 770 000 | 12 390 000 | 420 000 |
| S | ppb | 165 000 | 12 000 | 2 570 | 120 |
|  | nM | 1 870 000 | 140 000 | 29 200 | 1 300 |
| <sup>56</sup>Fe | ppb | 500 | 240 | 1 170 | 140 |
|  | nM | 9 000 | 4 400 | 20 800 | 2 400 |
| P | ppb | 1 100 | 1 300 | 630 | 630 |
|  | nM | 35 000 | 42 000 | 20 000 | 20 000 |
| Zn | ppb | 320 | 70 | 303 | 27 |
|  | nM | 4 900 | 1 100 | 4 600 | 400 |
| <sup>88</sup>Sr | ppb | 7 870 | 440 | 246 | 16 |
|  | nM | 89 400 | 5 000 | 2 800 | 180 |
| Ba | ppb | 638 | 21 | 291 | 17 |
|  | nM | 4 590 | 150 | 2 100 | 120 |
| Ni | ppb | 101 | 4 | 95 | 4 |
|  | nM | 1 680 | 73 | 1 585 | 61 |
| Mn | ppb | 46 | 2 | 79 | 8 |
|  | nM | 832 | 31 | 1 430 | 140 |
| Cr | ppb | 50 | 3 | 49 | 4 |
|  | nM | 956 | 66 | 948 | 76 |
| Cu | ppb | 88 | 58 | 55 | 10 |
|  | nM | 1 390 | 930 | 870 | 160 |

\* – used for culturing cells in Ba<sup>2+</sup>-loading experiments and as a rinsing solution for preparation of ICP-MS samples

## Notes

### Competing Interest Statement

The authors have declared no competing interest.

